# A Novel Process Maintaining Glycerophospholipid Homeostasis in Mammalian Cells

**DOI:** 10.1101/841221

**Authors:** Martin Hermansson, Satu Hänninen, Matti Kjellberg, Pentti Somerharju

## Abstract

Glycerophospholipid (GPL) homeostasis in eukaryotic cells is thought to be maintained via biosynthesis, degradation and acyl chain remodeling. Here we provide evidence for an additional process termed “head-group remodeling” where other GPLs, when in excess, are rapidly converted to phosphatidylcholine and triacylglycerol. Mass spectrometric studies showed the formation of diacylglycerol, but not phosphatidic acid, from the exogenous GPL thus indicating that the first step is catalyzed by a phospholipase C-type enzyme. Consistently, triacylglycerol formation was significantly inhibited by the knock-down of several PLCs, but not phospholipase Ds. Second, we found that each exogenous GPL strongly inhibited the synthesis of the corresponding endogenous GPL class. Based on these and previous data we hypothesize how mammalian cells could coordinate the multiple processes contributing to GPL homeostasis in mammalian cells. In conclusion, this study provides the first evidence that head group remodeling plays an important role in GPL homeostasis in mammalian cells.

## Introduction

The membranes of mammalian cells contain 7 major glycerophospholipid (GPL) classes, i.e. phosphatidylcholine (PC), -ethanolamine (PE), -inositol (PI), -serine (PS), -glycerol (PG), phosphatidic acid (PA) and cardiolipin (CL). Remarkably, the relative concentrations of these GPLs are kept within close limits in mammalian cells (1), obviously because deviations from the “optimal” composition can have dire consequences (2–6).

The well-known processes maintaining GPL homeostasis in mammalian cells are (*i*) biosynthesis, (*ii*) degradation and (*iii*) acyl chain remodeling. Among these, biosynthesis and degradation in particular must be accurately coordinated because competition between these opposing processes would be energetically costly. Indeed, compelling evidence for strict coordination of synthesis and degradation comes from studies in which the synthesis of PC, PE or PS was boosted by overexpressing the rate limiting synthetic enzyme ((7–11). Conversely, inhibition of the synthesis of those GPLs diminished their degradation in proportion ((12–15)). Surprisingly, however, nothing is known how biosynthesis and degradation are mechanistically coordinated except for the obvious fact that the coordination must rely on factors that (*i*) accurately “sense” the lipid concentration of the membranes and (*ii*) swiftly rely the information to synthetizing and degrading enzymes.

Here, we provide strong evidence that beside biosynthesis and degradation “head-group remodeling”, an previously unconsidered process, most likely plays an important role in GPL class homeostasis in mammalian cells. In addition, we show here that all exogenous GPLs strongly inhibit the synthesis of the corresponding endogenous GPL class, thus strongly suggesting that feed-back inhibition is not limited to some GPLs only, as shown previously, but is a universal mechanism in the regulation of GPL biosynthesis. Yet, we found that increasing the concentration of negatively charged GPLs in the cells stimulate the synthesis of PC apparently by increasing the activity of cytidylyltransferase (CCT). Based on these and previously published data we hypothesize how mammalian cells could coordinate the multiple processes contributing to GPL homeostasis.

## Results

### Introduction of exogenous GPLs to cells

In order to load a particular GPL (PC, PE, PS, PI, PG or PA) the cells were incubated in a medium containing small vesicles (100-200 nmol) consisting of a di-16:1-GPL, POPC and cholesterol in the presence of 1-2 mM methyl-β-cyclodextrin (mβ-CD). This protocol allows for efficient introduction of specific GPLs to cells without apparent adverse effects, such as extraction of cholesterol or PC from cells or compromised viability (16). Exogenous GPL species with two 16:1 acyl chains (di-16:1-GPLs) were used because they (*i*) are avidly transferred by mβ-CD and (*ii*) are very minor species in native HeLa cells (see legend of Fig. 1), which greatly simplifies monitoring the conversion of the loaded GPL to other lipids by mass spectrometry. No apparent adverse effects on cell viability or morphology were observed upon loading of the exogenous GPLs to the cells consistent with previous data (16). The loading method used allowed us to increase the cellular concentration of the acidic GPLs up to several-fold and that of PC and PE by 10-30%.

**Figure 1.**
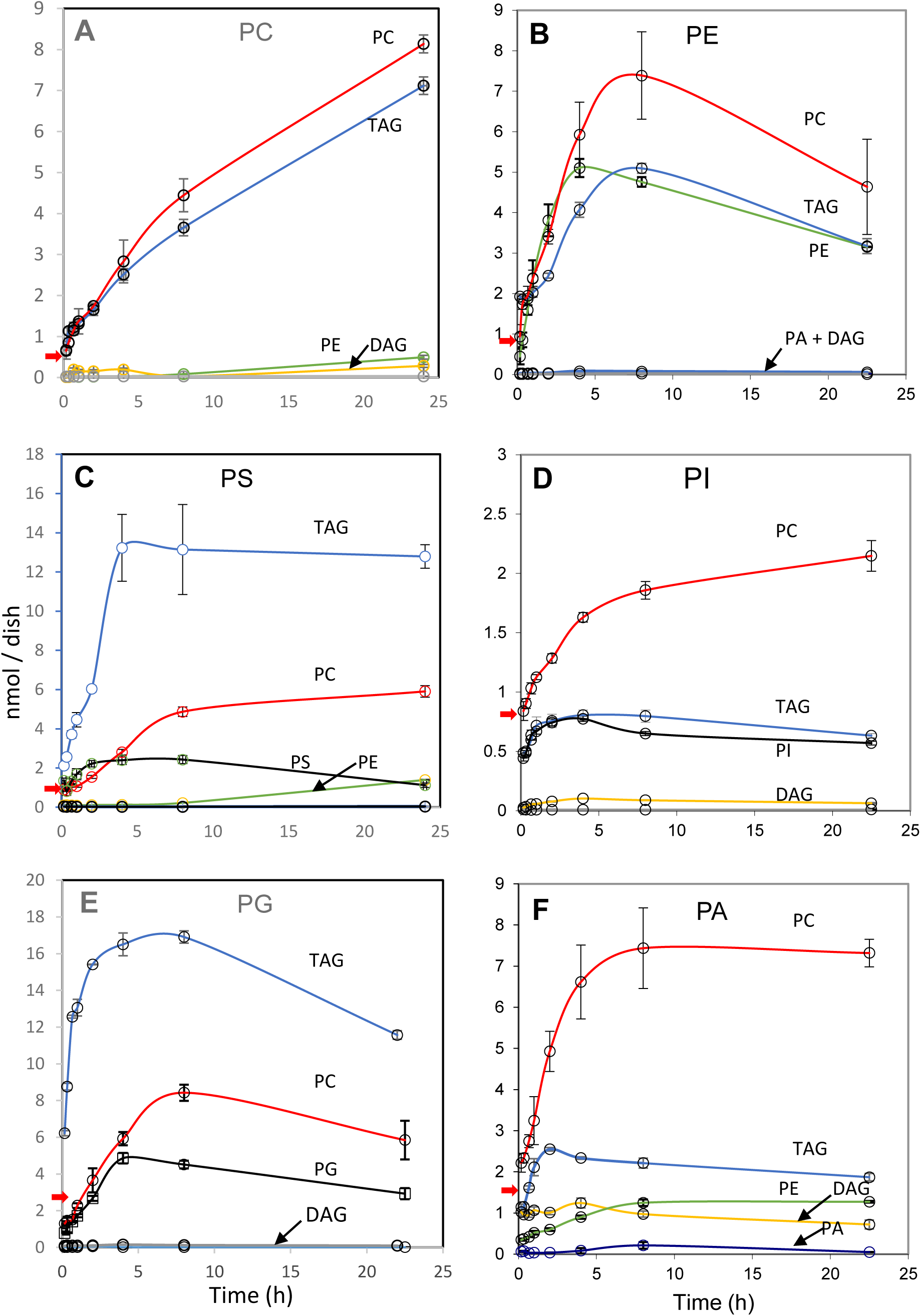
Conversion of exogenous GPLs to other lipids. Hela cells were incubated in a medium with vesicles containing either di-16:1-PE (A), -PS (B), -PI (C), -PG (D) or -PA (E) and 1 mM mβCD for the indicated time and then the lipids were extracted, methylated and analyzed with mass spectrometry as detailed under Materials and Methods. The lipids analyzed are indicated by abbreviations at the curves. Error bars are two small in some cases to be clearly visible. Red arrow at the Y-axis of the plots indicates the amount of endogenous di-16:1-PC in HeLa cells. The sum of endogenous TAGs containing two or three 16:1 acyl chains was 1.7 ±1 nmol (n = 2), while the di-16:1 species was negligible in the other GPLs and DAG (<0.5 % of total lipid in the class). The data are mean ± S.D. (n = 3). The experiment was repeated several times under similar conditions with parallel results.

### Exogenous GPLs are rapidly converted to PC and TAG

Remarkably, each exogenous GPL tested was rapidly converted to PC and TAG (Fig. 1) containing an intact di-16:2-DAG moiety derived from the precursor. This was demonstrated by experiments where Triacsin C, a potent inhibitor of acyl-CoA formation (17), was included in the medium. Triacsin C had no effect of the conversion of di-16:1-PE to di-16:1-PC (Fig. 2A), thus excluding the possibility that release of 16:1 fatty acid from the precursor and its subsequent reincorporation via de-novo synthesis is involved in the formation of di-16:1-PC. As expected, Triacsin C nearly abolished the conversion of the exogenous di-16:1-PE to di-16:1-TAGs (= TAGs containing 2 - 3 16:1 chains; Fig. 2A, Fig. S1), which requires acyl-CoAs for the addition of an acyl chain to di-16:1-DAG. Parallel results were obtained for PG (Fig. S1) and di-16:1-PS -and -PA (data not shown). The details of conversion of the different di-16:1-GPLs to other lipids were as follows.

**Figure 2.**
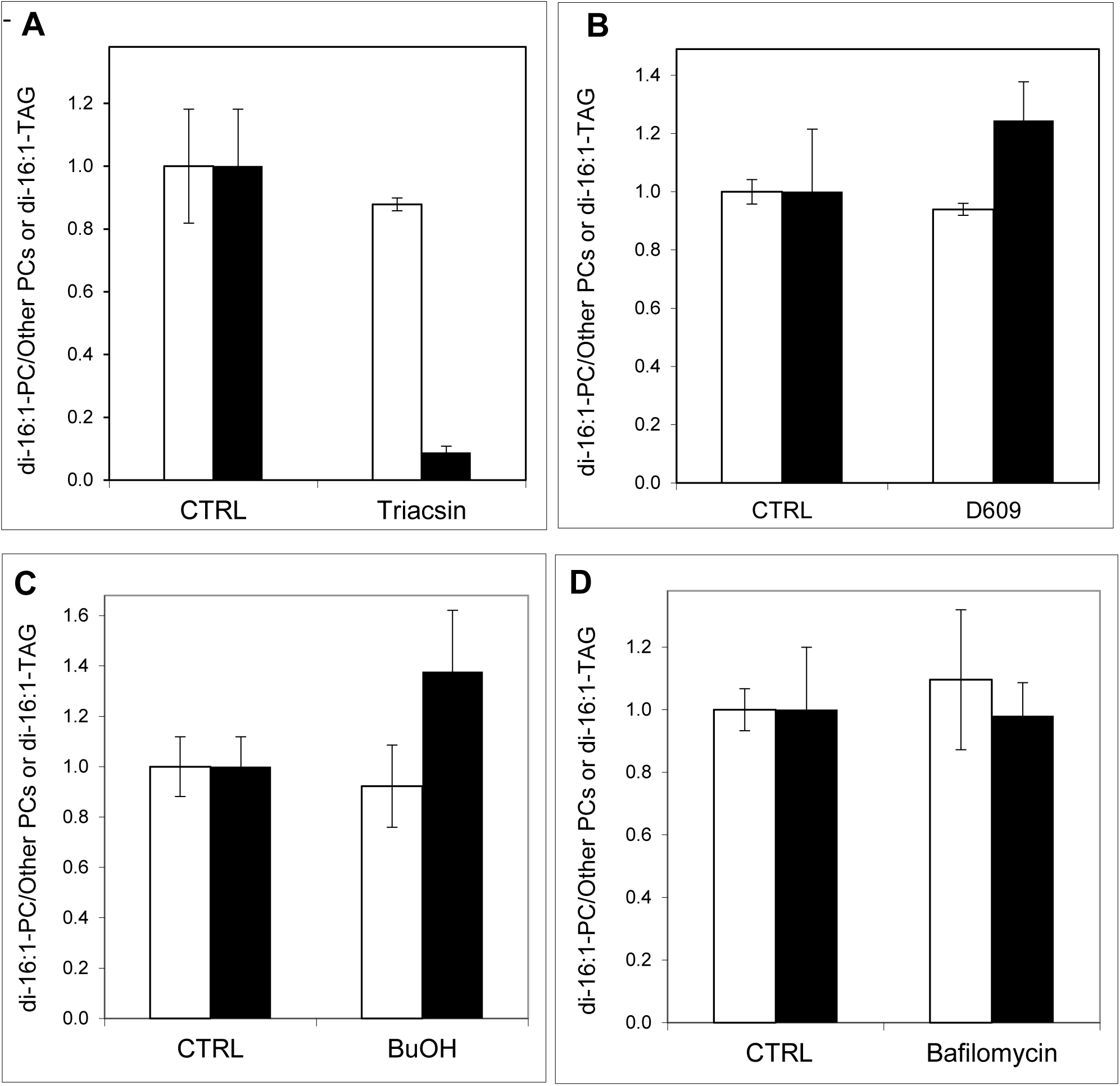
Effect of Triacsin C, D609, 1-Butanol or Bafilomycin on the conversion of di-16:1-PE to PC or TAG. HeLa cell monolayers were incubated with 20 µM Triacsin C (A), 50 µg/ml of D609 (B), 50 mM 1-butanol (C) or 0.033 µg/ml of Bafilomycin (D) for 1 h and then di-16:1-PE was loaded to cells for 3-4 h and its conversion to di-16:1-PC (white columns) and -TAG (black columns) was determined as detailed under Materials and Methods for the variation of cell number/dish. The data were normalized and are means ± standard deviation (n = 3). Each experiment was repeated was once or twice with similar results.

### PC

A significant fraction of exogenous di-16:1-PC was rapidly converted di-16:1-TAG (Fig. 1A) and an early peak of di-16:1-DAG was observed (Fig. S2A), suggesting that the PC was first hydrolyzed by a PLC (or a similar enzyme) to DAG which was then acylated to TAG. Consistently, only minor and invariable amounts of di-16:1-PA was observed (Fig. S2A).

### PE

Also exogenous di-16:1-PE was rapidly converted to di-16:1-PC (≈7 nmol at 8 h) and -TG (≈5 nmol at 8 h; Fig. 1B). Notably, again much more of di-16:1-DAG than di-16:1-PA was observed at any time (Fig. S2B).

### PI

Di-16:1-PI incorporated to the cells somewhat less efficiently than the other GPLs but was also rapidly converted to PC and TAG (Fig. 1D). A major increase of di-16:1-DAG but no increase of Di-16:1-PA was observed (Fig. S2C).

### PS

Since PS is rapidly decarboxylated to PE in mammalian cells, 1 mM hydroxylamine, which blocks PS decarboxylase (18) was added to the cell medium 1 h before the addition of the di-16:1-PS-containing vesicles. Again large amounts of di-16:1-TAG (≈13 nmol) and -PC (≈5 nmol) were are formed by 8 h of incubation (Fig. 1C). In contrast to PC and PE, there was no obvious increase of di-16:1-DAG (Fig. S2D), and since di-16:1-PA could not be determined in these cells (see Materials and Methods), these data neither indicate nor exclude that di-16:1-PS was hydrolyzed by a PLC.

### PG

Di-16:1-PG was very rapidly and effectively converted to TAG (≈16 nmol at 8 h) and PC (≈8 nmol at 8 h) (Fig. 1E). Again, di-16:1-DAG was much more abundant than di-16:1-PA at any time studied (Fig. S2E).

### PA

Di-16:1-PA was rapidly and effectively converted to di-16:1-PC (≈7 nmol at 7 h) and also significant amounts of TAG was formed (Fig. 1F). In a notable contrast to the other GPLs tested, significant amounts of di-16:1-DAG was present already at 10 min (Fig. S2F), indicating that di-16:1-PA was very rapidly dephosphorylated, presumably by a lipid phosphatase acting on the plasma membrane (19).

### Involvement of PLCs vs. PLDs in the conversion

The data in Fig. S1 is consistent with the notion that exogenous GPLs are first hydrolyzed to DAG by a PLC or a similar enzyme. To study this, we first tested the effect of D609, a PLC inhibitor (20). However, this compound had no effect on the formation of di-16:1-PC or -TAG from exogenous PE (Fig. 2B) or PS and PI (Fig. S6). Since D609 may not inhibit all cellular PLCs and inhibits sphingomyelin synthases as well (20), we knocked down the PLCs significantly expressed in HeLa cells, i.e., PLCB1, -B3, -D3, -E1, -G1, -H2 and - L2. The knock down of each PLC had a significant effect the formation of TAG from di-16:1-PE while the knock-down of the PLDs had no effect (Fig. 3A). Parallel results were obtained for exogenous do-16:1-PI and, to a lesser degree, for di-16:1-PS and -PG (Fig. S3). On the other hand, the knock down of PLCs had no effect on the conversion of di-16:1-PE to PC (Fig. 3B), while a modest reduction in the conversion of di-16:1-PI, -PS and -PG to PC was observed upon the knock-down of several PLCs (Fig. S4). The reason for the differences between the di-16:1-GPL precursors in the conversion to TAG vs. PC in the PLC knock down cells are not clear at this time. As found for PE, the knock-down of PLDs had little effect on the formation of di-16:1-TAG or -PC from other GPLs (Figs. S3 and S4).

**Figure 3.**
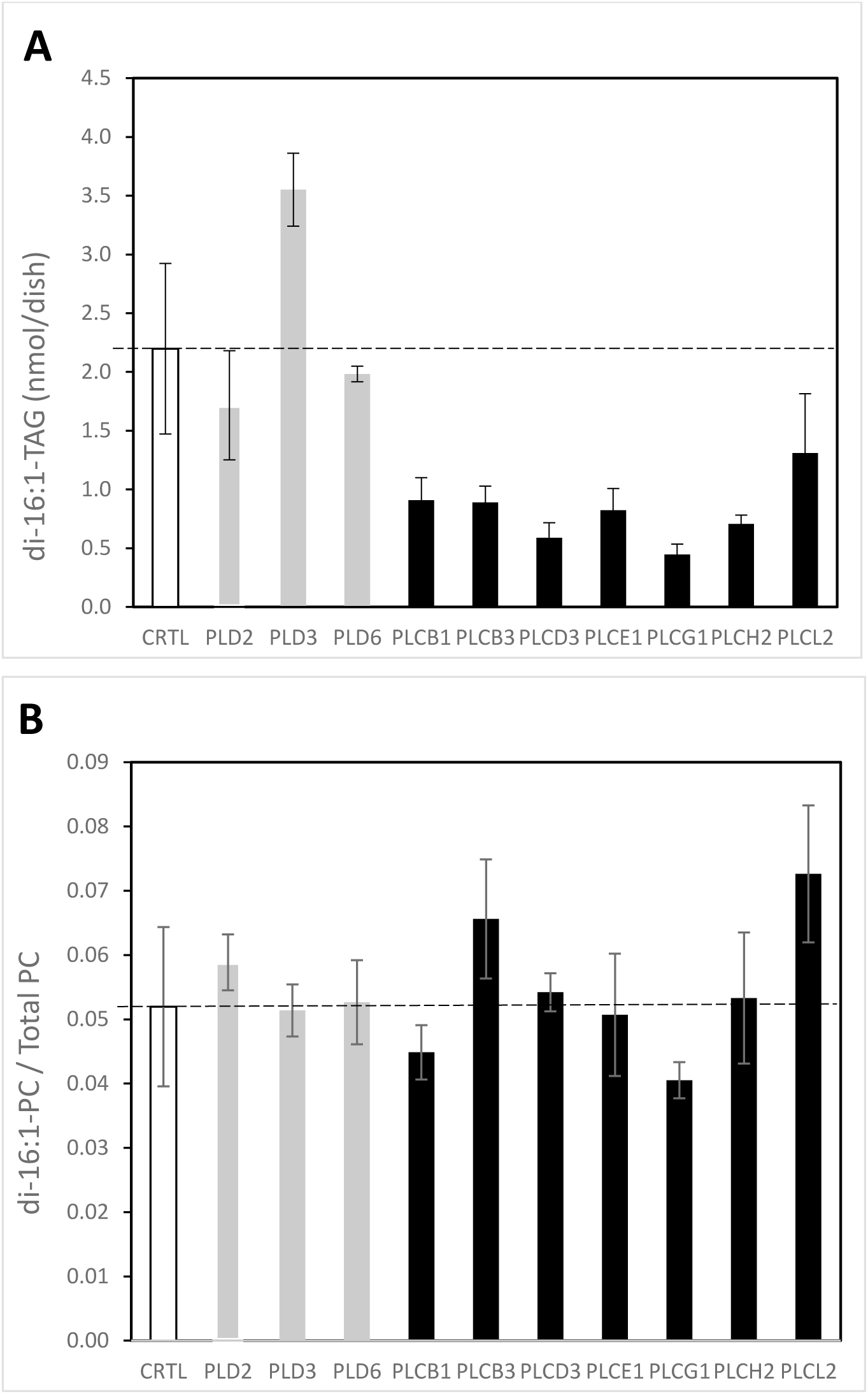
Effect of PLC or PLD knock-down of the conversion of exogenous di-16:1-PE to TAG (A) or PC (B). The indicated PLC or PLD was depleted by transfecting HeLa cells with the corresponding siRNA or CTRL siRNA for 72 h and then di-16:1-PE was loaded to the cells as specified under Materials and Methods. After 7 h the cells were washed and the lipids were extracted, methylated and analyzed with mass-spectrometry. For other details see the legend of Fig. 2. Data are means ± S.D. (n = 3)

To confirm that PLC-type enzymes rather than PLDs are catalyzing the first, committed step of the head group remodeling of exogenous GPLs, we tested the effect of 1-butanol known to prevent the formation of PA by PLDs and forming phosphatidylbutanol instead (21). As shown in Fig. 2C, 1-butanol had no significant effect on the formation of TAG or PC from PE or PS consistent with the notion that PLDs are not involved in the conversion.

### Involvement of other GPL metabolizing enzymes

Theoretically, exogenous PS could be converted to PC by PSS1 or PSS2 acting in reverse (22). Thus we knocked down both of these enzymes simultaneously so that only 6 ± 1.4 (n=2) % of PSS1 and 15 ± 3.5 (n = 2) % of PSS2 mRNA remained (Table S2). No significant effect on the conversion of exogenous PS to PC or TAG was observed (Fig. S5). Exogenous PE could be converted to PC by CEPT or EPT acting in reverse followed by conversion of the DAG formed to PC by CPT or CEPT (1). However, the knock down of any of these enzymes by ≈90% has an insignificant effect of the conversion of exogenous PE to PC (Fig. S6). Yet, exogenous PE could be converted to PC by the PE methyl transferase (PEMT). However, depletion of PEMT messenger in HeLa cells by 95% (Table S2) had no detectable effect on the conversion of exogenous PE to PC (Fig. S7).

In mammalian cells DAG is also formed from PC in a reaction catalyzed by SM synthase 1 or 2 (23). To study if either of these enzymes is involved in the conversion of exogenous PC to DAG, we knocked them down by siRNAs and then studied the conversion of exogenous di-16:1-PC to TAG. Indeed, the knock down of either enzyme significantly reduced TAG formation and the KD of SMS2 had a larger effect (≈80% reduction; Fig. 4), suggesting that the conversion of exogenous PC to DAG mainly occurs at the plasma membrane where SMS2 is located (23). The fact that D609 significantly inhibited the conversion of PC to TAG (data not shown) is consistent with the involvement of SMSs since these enzymes are inhibited by D609 (20).

**Fig. 4.**
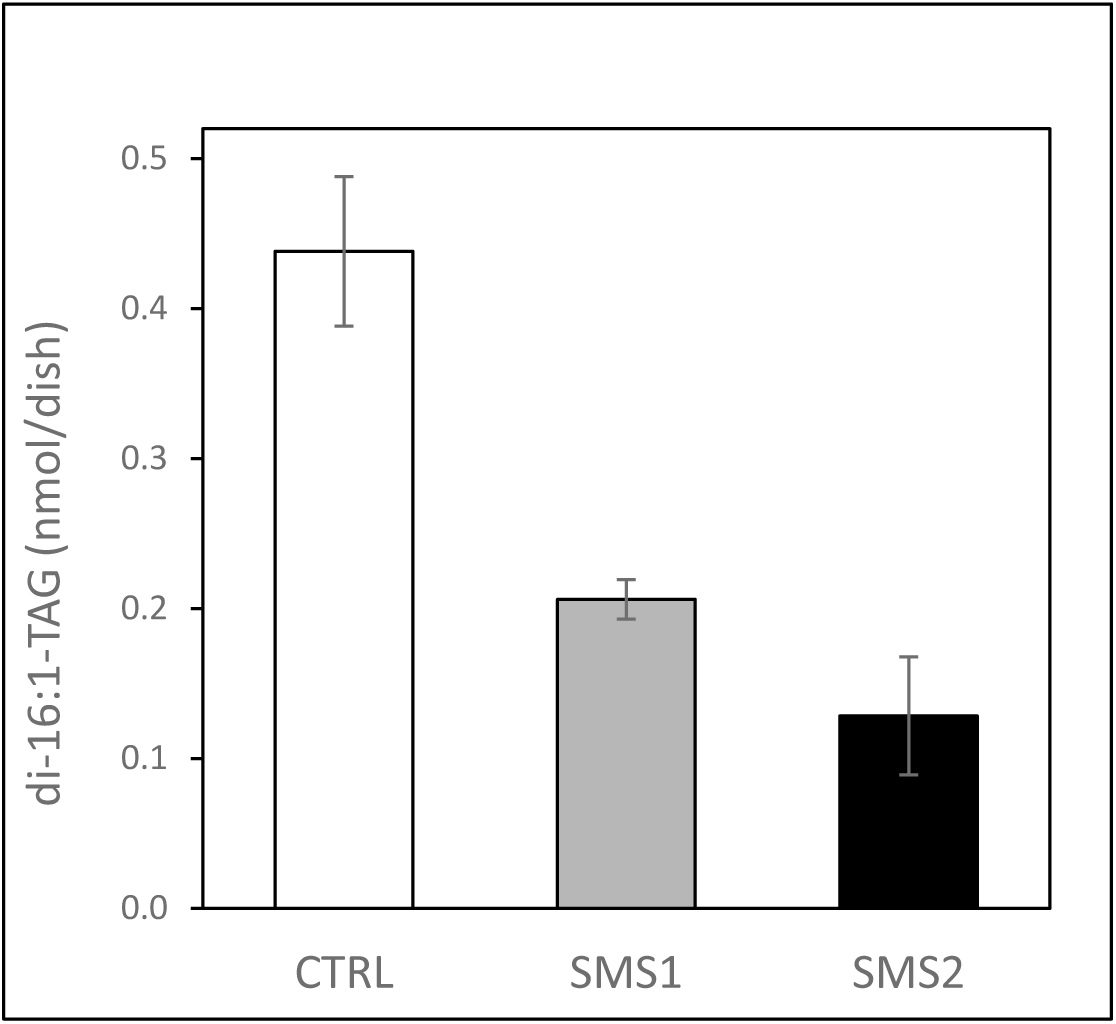
Effect of SMS1 or SMS2 knock-down on the conversion of di-16:1-PC to TAG. Either SMS1 (gray column) or SMS2 (black column) was depleted in HeLa cells and the conversion of exogenous di-16:1-PC to di-16:1-TAG was monitored as indicated in the legend of Fig. 5. White column: cells treated with a nonspecific (AllStars) siRNA. For other details see the legend of Fig. 2. Data are means ± S.D. (n = 3).

Exogenous PA could be converted to DAG by a lipid phosphate phosphatase (LPP) present a the plasma membrane of mammalian cells (19) and therefore we knocked down LPP1, −2 and −3 in HeLa cells. As shown in Fig. 5A, the knock down of LPP1 and LPP3 had a significant effect of the conversion of exogenous PA to TAG while that of LPP2 had no effect. No effect on the conversion of exogenous PA to di-16:1-PC was observed for any LPP knock down cells (Fig. 5B). We conclude that LPP1 and LPP3 are involved in the conversion of exogenous PA to DAG as indicated by di-16:1-TAG formation. The lack of effect by the knock down on the conversion of PA to PC is probably again explained by that DAG is not rate-limiting in PC synthesis.

**Figure 5.**
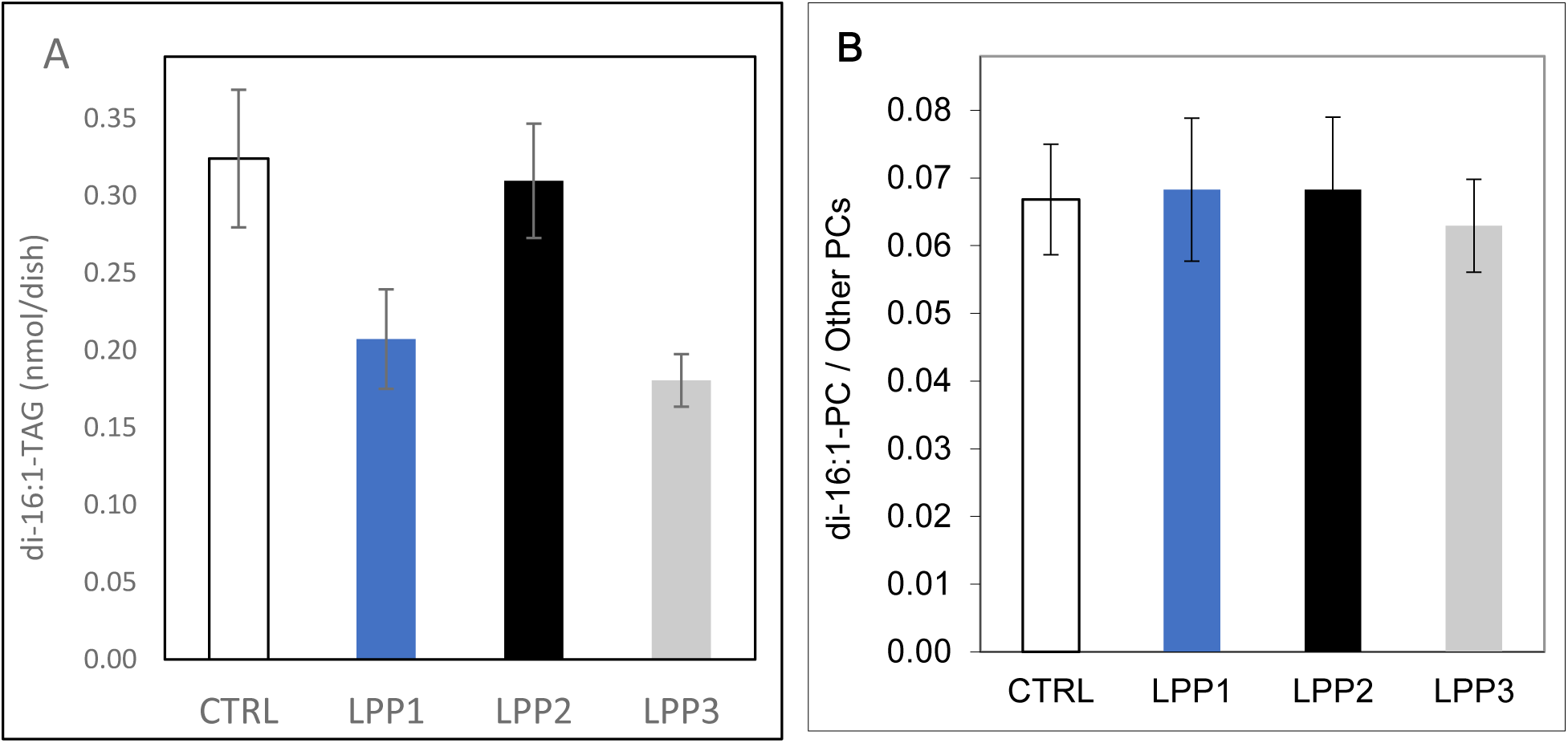
Effect of lipid phosphatase knock-down on the conversion of di-16:1-PA to TAG. LPP1, −2 or −3 was depleted in HeLa cells and the effect on the conversion of exogenous di-16:1-PA to TAG (A) or PC (B) was determined as detailed under Materials and Methods.

To test the possible involvement of putative base-exchange enzymes, we investigated whether the conversion of exogenous PE or PS to PC takes place in the absence on an intact CDP-choline pathway. To this end, we utilized CHO-58 cells that contain a heat-sensitive CTα, the rate-limiting enzyme in PC synthesis (24). At 33 ºC the CHO-58 cells grow similarly to wild-type cells and have a normal GPL composition while at 40 ºC, where CT is inactive, the cells cannot synthetize PC (25, 26)which leads to apoptosis (27). We found that at 33 ºC the CHO-58 cells took up exogenous PS or PE and converted them to PC and TAG like the wild-type CHO-K1 cells. However, at 40 ºC no conversion of exogenous PE or PS to PC was observed, whereas di-16:1-TAG still accumulated (data not shown). These data demonstrate that the conversion of exogenous PE or PS to PC requires an intact CDP-choline pathway thus excluding the involvement of sc. base-exchange mechanism.

Finally, we considered the possibility that a putative lysosomal PLC, found previously in rat liver (28), is involved in the conversion of exogenous GPLs to PC in the present study. However, blocking the acidification of lysosomes and endosomes with Bafilomycin A1 (26, 29)had little or no effect on the conversion of exogenous PE to PC or TAG (Fig. 2D). Consistent data were obtained for exogenous di-16:1-PS and -PG (data not shown). Thus the involvement of putative lysosomal PLCs in the conversion of exogenous GPLs to PC or TAG is unlikely.

### Exogenous GPLs markedly inhibit the synthesis of corresponding endogenous GPL

It has been demonstrated previously that exogenous PS effectively inhibits the synthesis of PS in CHO cells (29, 30). Similar albeit less detailed data has been obtained for exogenous PI in rat pituitary cell membranes (31), and exogenous LPC and a PC analogue inhibited the synthesis of PC in a murine macrophage cell line (32). Here we observed that in fact each exogenous GPL studied inhibited the synthesis of the corresponding endogenous GPL (Fig. 6). The inhibition was particularly strong for PI, PG and PS and modest for PC and PE. The fact that less inhibition was observed for PE and PC is most likely due to that loading protocol used allowed us to increase the mole fractions of PC and PE (relatively) far less as that of the other GPLs, simply because PE and PC and are 10 to 20 times abundant than the other GPLs in the cells (16).

**Figure 6.**
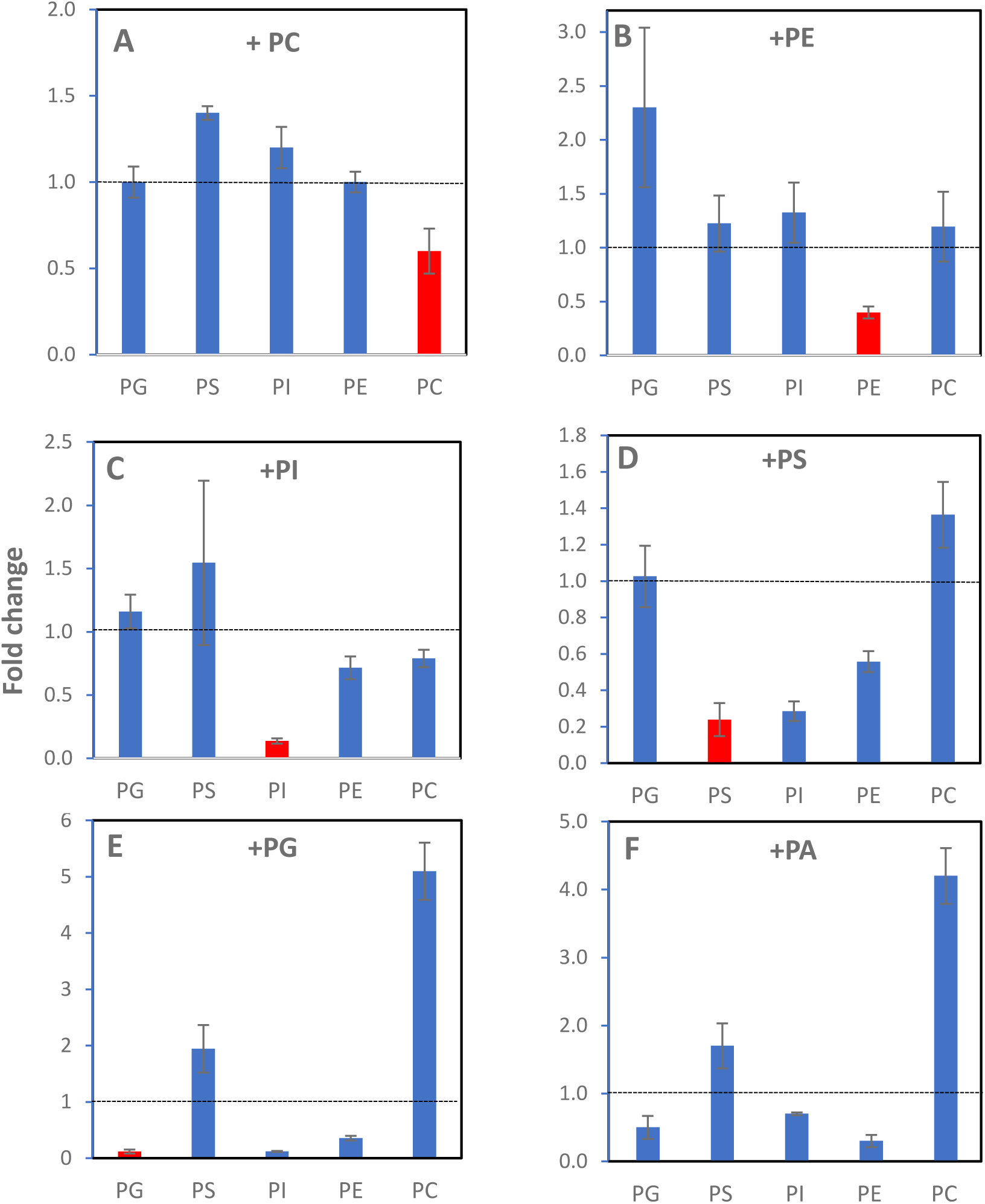
Effect of loading of exogenous GPLs on GPL synthesis. HeLa cells were incubated for 1 hour in a medium with vesicles containing di-16:1-PC (A), -PE (B), -PS (C), -PI (D) or -PG (E) and 1-2 mM mβCD for an hour and then a mixture of D_9_-choline, D_4_-ethanolamine, D_3_-serine, D_6_-inositol and ^13^C_3_-glycerol was added. After 6 h at 37 ºC the cellular lipids were extracted, methylated and analyzed by mass spectrometry (see Materials and Methods). When PS was loaded to the cells (panel D) the medium contained 1 mM hydroxylamine which prevents the decarboxylation of PS to PE. The GPL loaded to the cells is indicated by a red column in each panel. The data are means ± S.D. (n = 6). There is no column for PA in panel F since the synthesis of this GPL could not be determined due to the lack of a selective MS/MS scan for PA. Dashed line indicates the level where there is no effect on synthesis. The experiment was repeated several times under similar conditions with analogous results.

For PC, PE and PI the inhibition was specific, i.e., only the synthesis of the corresponding GPL was significantly affected. In contrast, beside inhibiting its own synthesis by ∼90%, exogenous PG also markedly inhibited the synthesis of PI by ∼90% and PE by ∼65%. Exogenous PS inhibited its own synthesis by ∼75% and that of PI by ∼70% and PE by ∼45%. Exogenous PA significantly inhibited the synthesis of PI, PG and PE. Notably, the inhibition of PE synthesis by PA could be direct or indirect i.e., due to the PE derived from exogenous PA (cf. Fig. 1F).

Intriguingly, exogenous PG and PA increased strongly (4 to 5-fold) and exogenous PS slightly (1.4-fold) the synthesis of PC (Fig. 6E,F). Exogenous PC had no effect on the synthesis of the other GPLs excluding PS whose synthesis was increased by 1.4 - 2 -fold (Fig. 6A). Loading of PC to the cells significantly increased the degradation of the endogenous PC (data not shown) consistent with previous studies where PC synthesis was boosted by overexpressing CCT (8, 9).

In summary, increasing the concentration of a negatively charged GPL (or PE) in the cells led to three distinct homeostatic responses: (i) rapid conversion of the exogenous GPL to PC and TAG, (ii) efficient inhibition of the synthesis of the corresponding endogenous GPL and (iii) stimulation of PC synthesis. Exogenous PC was effectively converted to TAG and inhibited its own synthesis.

## Discussion

### GPL head group remodeling is a novel homeostatic process

Here we provide the first evidence that in mammalian cells PE, PS, PI, PG and PA can be efficiently converted to PC and TAG in a process involving “head group remodeling”. Exogenous di-16:1-PC was effectively converted to TAG consistent with previous data on primary hepatocytes (33) but not significantly to other GPLs.

To study the mechanisms of the conversion, we inhibited or knocked down the GPL metabolizing enzymes potentially involved in this process, including all PLCs and PLDs significantly expressed in HeLa cells, PS synthases 1 and 2, CEPT, ETP, CPT, PEMT, SMS 1+2 and LPP1-3. Among these, only the knock down of PLCs significantly inhibited the formation of TAG from exogenous PE. Intriguingly, the knock-down of any PLC alone had a significant effect on the formation of TAG from PE implying that multiple PLCs could be involved in the conversion. It is intriguing that the knock down of the PLCs reduced the conversion of PE (Fig. 3) and PG (Fig. S3) to TAG since PLCs are thought to be specific for PIP2 or PC. However, to the best of out knowledge, the substrate specificities of the PLCs implicated here have not been thoroughly studied in cells. Alternatively, the PLCs may not act directly on the exogenous GPLs but rather regulate putative PLC-type enzyme(s) that carry out the hydrolysis. Clearly, further studies are needed to resolve the role of PLCs in head group remodeling of GPLs.

In contrast to conversion of PE to TAG, the knock down of PLCs had little or no effect on the conversion of exogenous PE to PC (Fig. 3) which may seem unexpected since DAG is the precursor of both PC and TAG. However, while the rate of TAG synthesis generally depends on the concentration of DAG, the rate-limiting step in PC synthesis is the synthesis of CDP-choline by CCT (34). Previous data also indicate that when DAG is abundant, it is preferentially used for the synthesis of TAGs rather than PC or PE (35). These phenomena probably explain why PLC knock down inhibited the conversion of di-16:1-PE to TAG but not to PC. Intriguingly, the knock down of some PLCs reduced the conversion of exogenous PS to PC but not to TAG (Fig. S3,4). Whether the differences in the metabolism of exogenous PE vs. PS are due to the more efficient incorporation of the latter to the cells (Fig. 1) remains to be studied.

The knock down of SMS1 or SMS2 significantly inhibited the conversion of exogenous PC to TAG (Fig. 4) which is to be expected as those enzymes produce DAG (and SM) from PC (23). The fact that the knock down of SMS2 had a greater effect than that of SMS1 suggests that much of the conversion takes place at the plasma membrane where SMS2 is mainly located.

In summary, while SMS1 and −2 seem to catalyze, at least in part, the conversion of exogenous PC to DAG and LPP1 and −3 to catalyze that of exogenous PA to DAG, PLCs are strongly implicated as the enzymes catalyzing (directly or indirectly) the initial step in the conversion of the other exogenous GPLs to DAG, the precursor of PC and TAG.

### Inhibition of biosynthesis by the end product is common to all major GPL classes

Previously Kuge and coworkers have shown that PS synthesis in Chinese hamster ovary cells cells in strongly inhibited by exogenous PS and that a specific arginine of PSS1 and PSS2 is critical for the inhibition (29, 30). Similar, albeit less detailed data were obtained for PI synthesis in early studies on rat pituitary membranes (31), and the synthesis of PC was shown to be inhibited by lyso-PC and a PC analogue (32). Here, we show that loading of any common GPL to cells strongly inhibits the synthesis of the corresponding GPL (Fig. 6). Accordingly, our data strongly suggest that inhibition of the synthesis by the end product is not restricted to some GPLs only but is a general mechanism of homeostatic regulation in mammalian cells.

Inhibition of biosynthesis by an exogenous GPL was specific in the case of PC, PE and PI, i.e. only the biosynthesis of the corresponding GPL was significantly inhibited (Fig. 6). In contrast, exogenous PG also markedly inhibited the synthesis of PI and PE and (less) that of PS. The inhibition of PI synthesis by PG is consistent with the previous findings showing that PI and PG are co-regulated, i.e., their sum concentration remains constant when the concentration of either GPL is altered (36, 37). The lack of inhibition of PG synthesis by exogenous PI could be due to the less efficient internalization of PI as compared to PG (cf. Fig. 1).

Beside its own synthesis, exogenous PS markedly inhibited the synthesis of PI and modestly that of PE. The negative charge of its head group could explain why PS inhibited the synthesis of PI since previous studies have shown that mammalian cells attempt to maintain the concentration of negatively charged GPLs in their membranes constant (29, 36). This could also be the reason why exogenous PA inhibited the synthesis of PG and PI (Fig. 6). On the other hand, the inhibition of PE biosynthesis by exogenous PA could result from that these GPLs have similar molecular shapes due to a small head group.

### Upregulation of PC synthesis by negatively charged GPLs

Loading of PS or PG to the cells upregulated the synthesis of PC (Fig. 6), most probably by activating CCT, the rate limiting enzyme of PC synthesis (34). Intriguingly, exogenous PI did not stimulate the synthesis of PC in contrast to PS and PG, possibly because PI is less potent in stimulating CCT activity that the other negatively charged GPLs, at least in vitro (38). The bigger and strongly hydrated inositol head group may shield the negative charge of the phosphate moiety thus hampering its interaction with positively charged residues of CCT which mediates its association with the membrane thus activating it (34). Another reason may be that less PI was internalized by the cells (cf. Fig. 1) and thus its concentration remained was not significantly increased in the nuclear or ER membranes where CCT acts (39).

### How are the different homeostatic processed coordinated?

As discussed in Introduction, it is commonly accepted that GPL class homeostasis in mammalian cells relies on balanced action of biosynthesis and degradation of the different GPLs. In this manuscript we provide strong evidence for an additional process, i.e. head group remodeling, which could play an important role in GPL homeostasis. Yet another process influencing the GPL composition of an individual organelle membrane is the translocation of GPLs to and from that membrane (40). Because of all those four processes can influence GPL composition and because there are so many GPL classes present in each membrane, it must be highly challenging to maintain GPL homeostasis in a membrane, i.e., it requires strict coordination of biosynthesis, degradation, head group remodeling and inter-organelle translocation. To date, the mechanism(s) underlying such coordination have not been uncovered, but we have recently hypothesized that the key coordinating factor is the *chemical activity* of the different GPL classes (41). According this hypothesis, abrupt composition-dependent changes in the chemical activity (≈ mole fraction) of the individual GPLs regulate all four homeostatic processes thus coordinating them to maintain homeostasis. Notably, chemical activity of cholesterol has been previously proposed to regulate its biosynthesis and intracellular translocation (42–45). Finally, we note that, beside the acute regulation discussed above there are also other, slower processes regulating GPL composition of mammalian cells operating under particular cellular states such as mitosis, differentiation and apoptosis (41).

## Materials and Methods

### Chemicals and reagents

Culture media and reagents and the solvents (LC-MS grade) were obtained from Fisher Scientific (Vantaa, Finland). D_9_-choline, D_3_-L-serine and D_4_-ethanolamine and ^13^C_3_-glycerol were from Cambridge Isotope Laboratories (Andover, MA), D_6_-myo-inositol from CDN Isotopes (Quebec, Canada), hydroxylamine (HA), methyl-β-cyclodextrin (mβ-CD), phospholipase D (*Streptomyces sp*ecies), formic acid, ammonium formate and ammonium hydroxide from Sigma. The primers were obtained from Metabion International AG. Phospholipid were purchased from Avanti Polar Lipids (Alabaster, AL) or were synthesized as previously (46, 47) and their purities were determined with thin layer chromatography and confirmed by mass spectrometry. Phospholipid concentrations were determined using a phosphate assay (48).

### Cell culture and RNA interference

HeLa cells were cultured in DMEM containing 10% fetal bovine serum, 100 units/ml penicillin and 100 μg/ml streptomycin under 5% CO_2_ in air at 37 °C. The cells were transfected with siRNAs (Table S1) using the Lipofectamine RNAiMAX reagent (Fisher Scientific) as described previously (49). The concentrations of the siRNAs in the cell medium and the knock-down efficiencies are indicated in Table S2.

### RNA isolation and RT-qPCR

RNA was extracted from cells with Machery-Nagel NucleoSpin RNA II Kit (Biotop Oy, Turku, Finland) and two µg was transcribed to cDNA using RevertAid reverse transcriptase (Thermo Scientific) and random hexameric primers. The qPCR reactions were carried out using 5 µl of cDNA (diluted 1:20) using specific primers (300 nM; Table S3) (Metabion International AG, Germany) in 20 µl of SsoAdvanced universal SYBR Green Supermix (Bio-Rad). Amplification was carried out on Bio-Rad C1000 Thermal Cycler by incubating for 15 s at 95 °C and then 45 s at 60 °C (35 cycles). The data were analyzed using the CFX Manager 3.1 software.

### Introduction of exogenous GPLs to cells

HeLa cells were grown to ∼ 80% confluency on 3 or 6 cm dishes. To load an exogenous GPL to cells 100 - 400 nmol of donor vesicles consisting of a di-16:1-GPL, POPC and cholesterol (1:1:2, mol/mol), mβ-CD (1 or 2 mM) were added to 1.5 ml of DMEM medium containing 2% FBS and the cells were kept in a cell incubator for up to 24 h at 37 °C.

### Effect of exogenous GPLs on GPL synthesis

Cells on 3 cm dishes were washed twice with PBS and then incubated in DMEM containing 1 - 2 mM mβCD and sonicated vesicles (see above) for 1 hour at 37 ºC in a cell incubator. Then D_9_-choline (100 μg/ml), D_4_-ethanolamine (100 μg/ml, adjusted to pH 7.0 by acetic acid), D_3_-L-serine (300 μg/ml) and D_6_-myo-inositol (100 μg/ml) and ^13^C_3_-glycerol (400 µg/ml) were added to the medium and the incubation was continued for up to 24 hours. The cells were then washed twice and the lipids were extracted and analyzed as indicated below. In the experiments with di-16:1-PS 1 mM hydroxylamine was included in the cell medium to prevent decarboxylation of PS to PE (18).

### Lipid analysis

Lipids were extracted from the cells using an acidic Folch method (50). After partitioning the lower phase was washed once with theoretical upper phase and then taken to dryness under nitrogen flow. The residue was dissolved in 1 ml of C/M (95:5), 5 µL of 0.5 M HCl, 50 µl of trimethylsilyldiazomethane (2 M) in hexane were added and the samples were incubated for 10 min at RT in order to methylate the phospholipids (51, 52). (*Note*: trimethylsilyldiazomethane is toxic and thus the procedure should be carried out in a fume hood while carrying personal safety equipment). Then 7 µL of acetic acid was added to quench the remaining methylation reagent. Notably, we observed that a significant fraction of PS is converted to methylated PA during the process, which precludes the analysis of the cellular PA species except for those insignificant in PS (e.g., di-16:1) The lipids were then re-extracted as above but without added acid, taken to dryness and dissolved in 50 µL of methanol and then analyzed by LC-MS on system consisting of Waters Acquity H-class UPLC system and Micro Premier triple quadrupole mass spectrometer. The column (Waters Acquity BEH C_18_, 1 x 100 mm) was eluted with a gradient of solvent A (acetonitrile/water (6:4, v/v) containing 10 mm NH_4_-formate and 1% NH_4_OH) to solvent B (2-propanol/acetonitrile (9:1, v/v) containing 10 mm NH_4_-formate and 1% NH_4_OH) as detailed previously (53). The methylated GPLs were detected using class-specific neutral-loss or precursor ion scanning in the positive ion mode (52). The following neutral losses were used for the unlabeled and head group - labeled GPLs, respectively: m/z 155, 159 (PE); m/z 213, 216 (PS); m/z 291, 297 (PI); m/z 203, 206 (PG) or m/z 143 (PA). Unlabeled and labeled PC were detected by scanning for the precursors of m/z of 198 or 207. Spectra were extracted from the chromatograms with the MassLynx software (Waters) and analyzed with the LIMSA software (54). DAG and TAG were detected using multiple reaction monitoring and were quantified using the QuanLynx software (Waters). Quantification of the lipids was based on the following internal standards: PC-40:2, PE-40:2, PS-28:2, PI-32:2, PG-40:2 and PA-40:2 and DAG-40:2 (for DAG and TAG).

### Other methods

Concentrations of the phospholipid stocks were determined using a phosphate assay (Bartlett 1970).

## Abbreviations

CL: cardiolipin
CCT: CTP:phosphocholine cytidylyltransferase
CEPT: choline/ethanolamine phosphotransferase
CHPT: choline cytidylyltransferase
DAG: diacylglycerol
EPT: ethanolamine phosphotransferase
GPL: glycerophospholipid
mβ-CD: methyl-β-cyclodextrin
PA: phosphatidic acid
PC: phosphatidylcholine
PG: phosphatidylglycerol
PE: phosphatidylethanolamine
PI: phosphatidylinositol
PS: phosphatidylserine
PEMT: phosphatidylethanolamine methyltransferase
PSS: phosphatidylserine synthase
PLA: phospholipase A
SMS: Sphingomyelin synthase
TAG: triacylglycerol

## Acknowledgements

We are grateful to Tarja Grundström for excellent technical assistance and to Academy of Finland and Sigrid Juselius Foundation for funding.

## Competing interests

The authors declare no competing interests.

