## Supplemental figures and tables for "A Novel Process Maintaining Glycerophospholipid Homeostasis in Mammalian Cells"

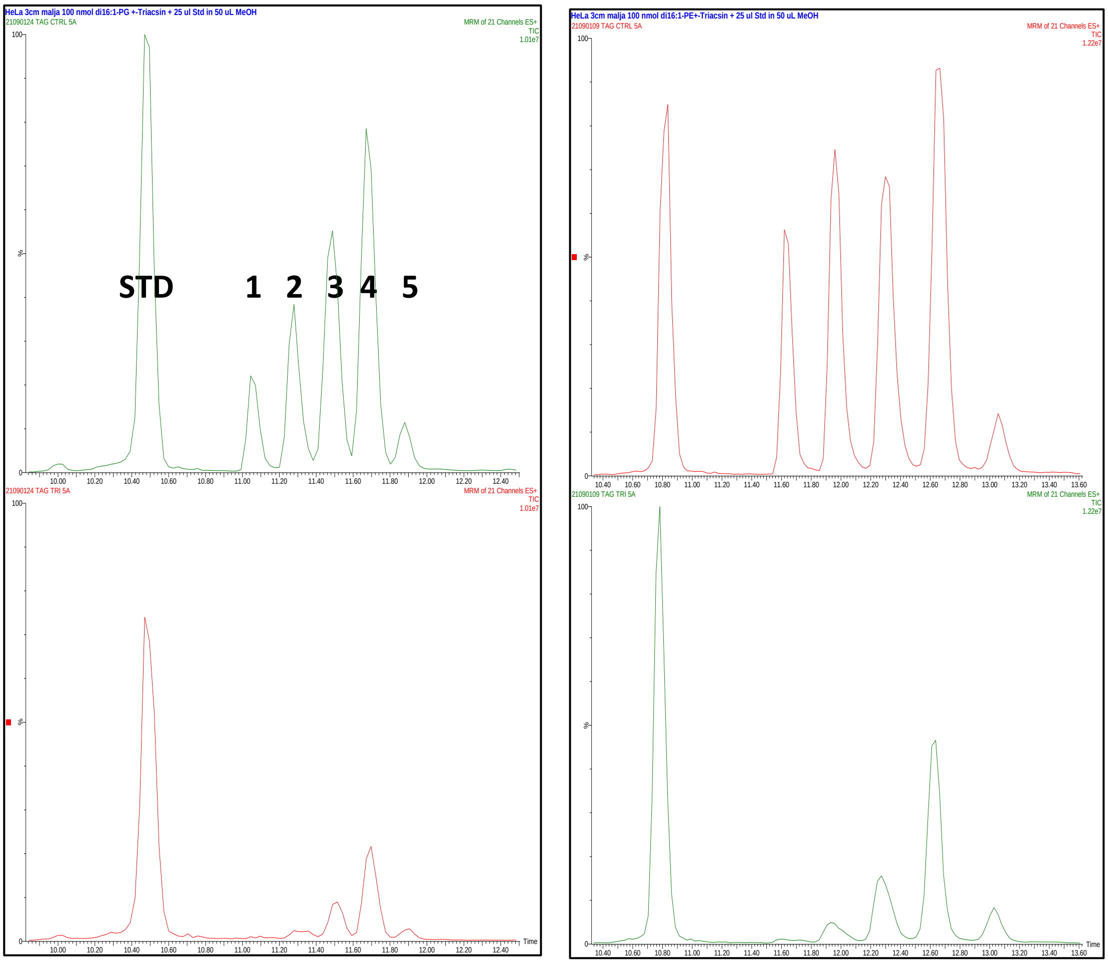


***Figure S1***. *LC-MS/MS chromatograms of TAG species from HeLa cells loaded with di-16:1-PE (left column) or di-16:1-PG (right column) in the absence (upper panels) or presence (lower panels( of Triacsin C for 5 h.* Triacsin C was added to medium 1 h before loading the cells with the exogenous GPL (see Materials and Methods). Identity of the peaks: STD = di-20:1-DAG standard; 1 = tri-16:1-TAG; 2 = 16:1/16:1/18:1-TAG; 3 = 16:1/18:1/18:1-TAG; 4 = 16:0/18:1/18:1-TAG; 5 = 18:0/18:1/18:1-TAG.

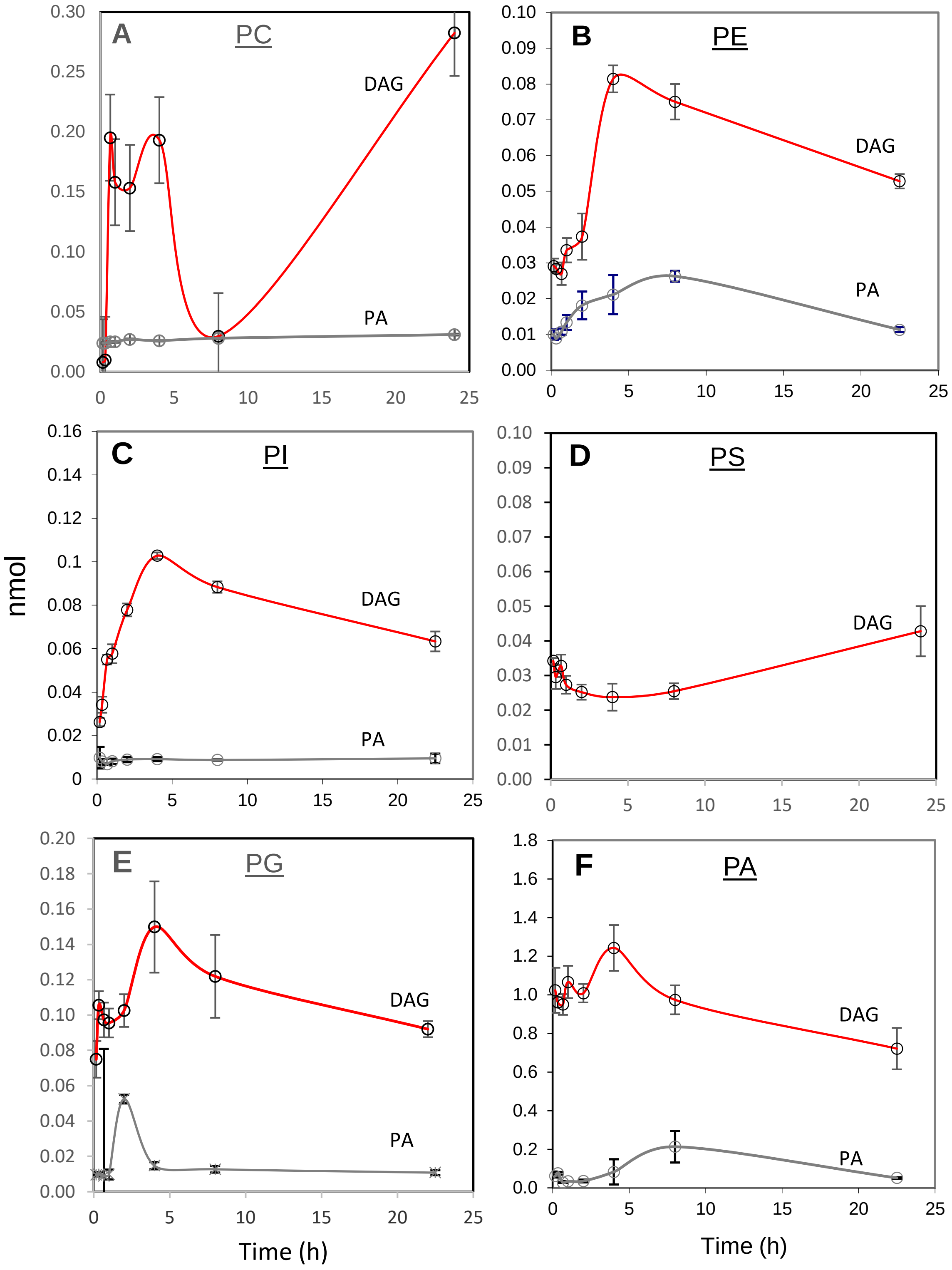


**Figure S2**. *Conversion of exogenous di-16:1-GPLs to di-16:1-DAG and -PA*. For details see the legend of Fig. 1 in the main text. The GPL loaded to cells is indicated at the top of each panel. Note that the Y-scale in the PA panel is much larger than in other panels and Di-16:1-PA could not be determined when PS was loaded to the cells (see Materials and Methods). Data are means ± S.D. (n = 3).

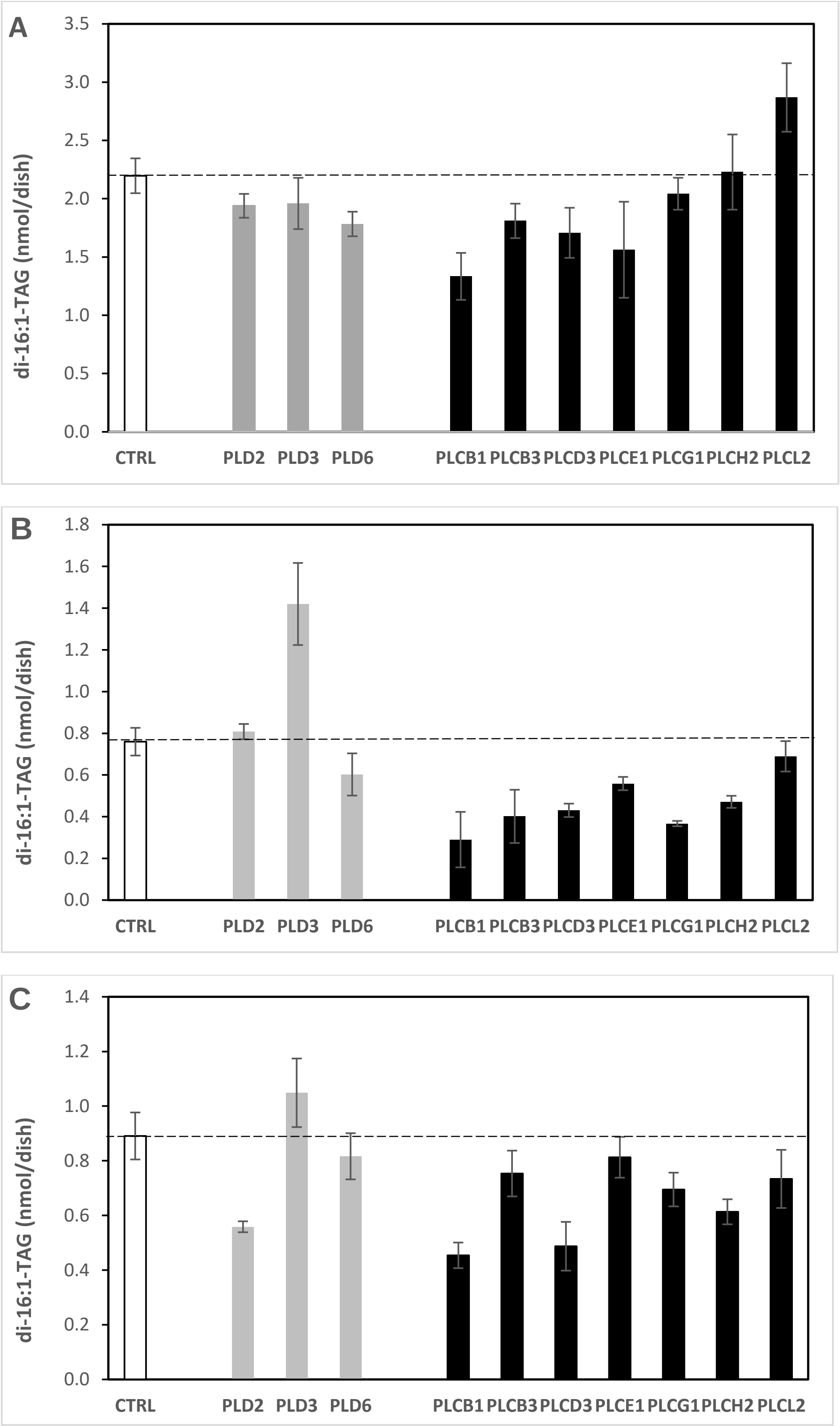


**Figure S3.** *Effect of PLC or PLD knock down on conversion of di-16:1-PS (A),-PI (B) or PG (C) to di-16:1-TAG***.** For details see Figure 4 in the main text. The data are mean ± S.D. (n = 3).

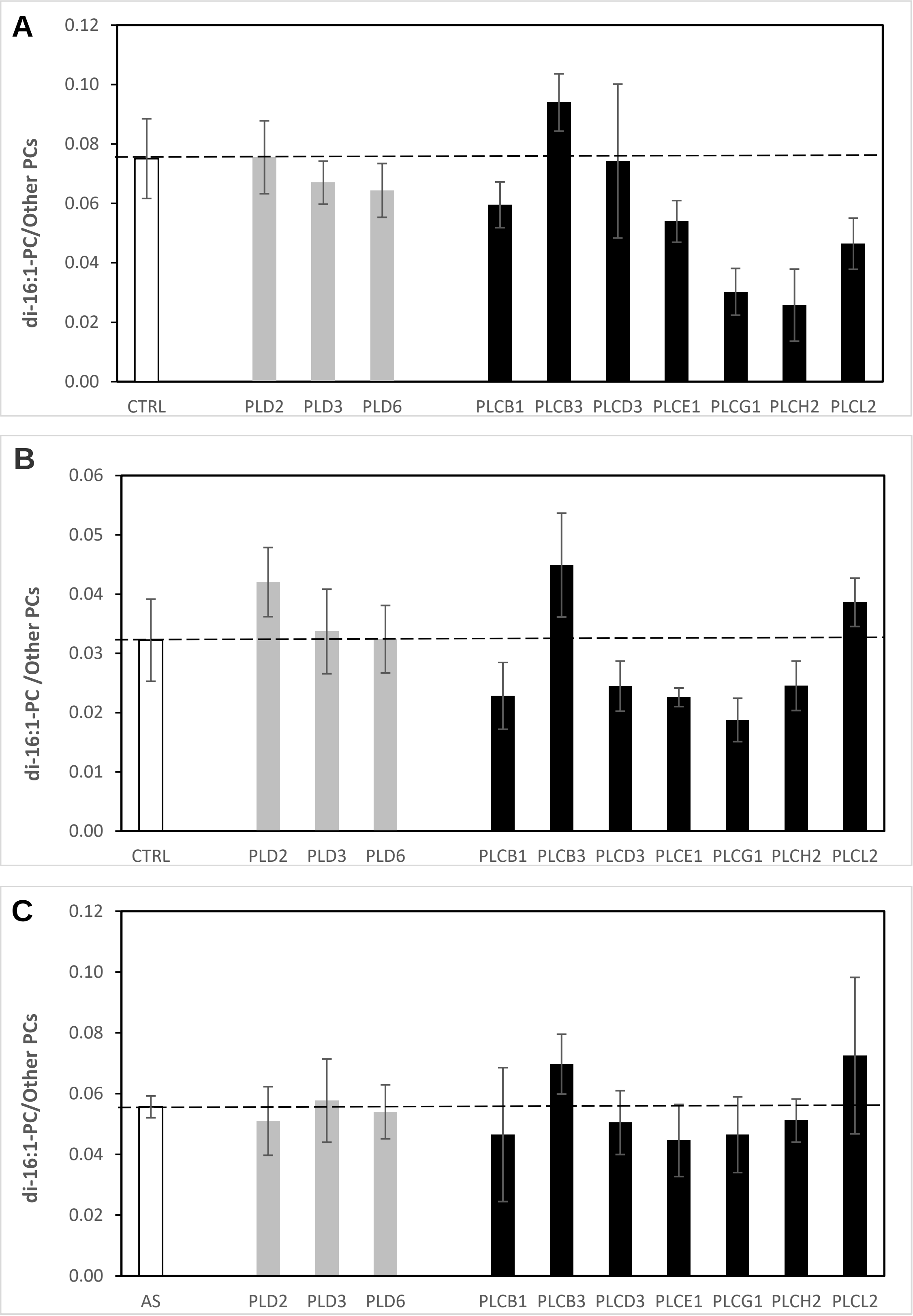


**Figure S4.** *Effect of PLC or PLD knock down on the conversion of di-16:1-PS (A), PI (B) or PG (C) to di-16:1-PC***.** For details see Figure 4 in the main text. The data are mean ± S.D. (*n* = 3).

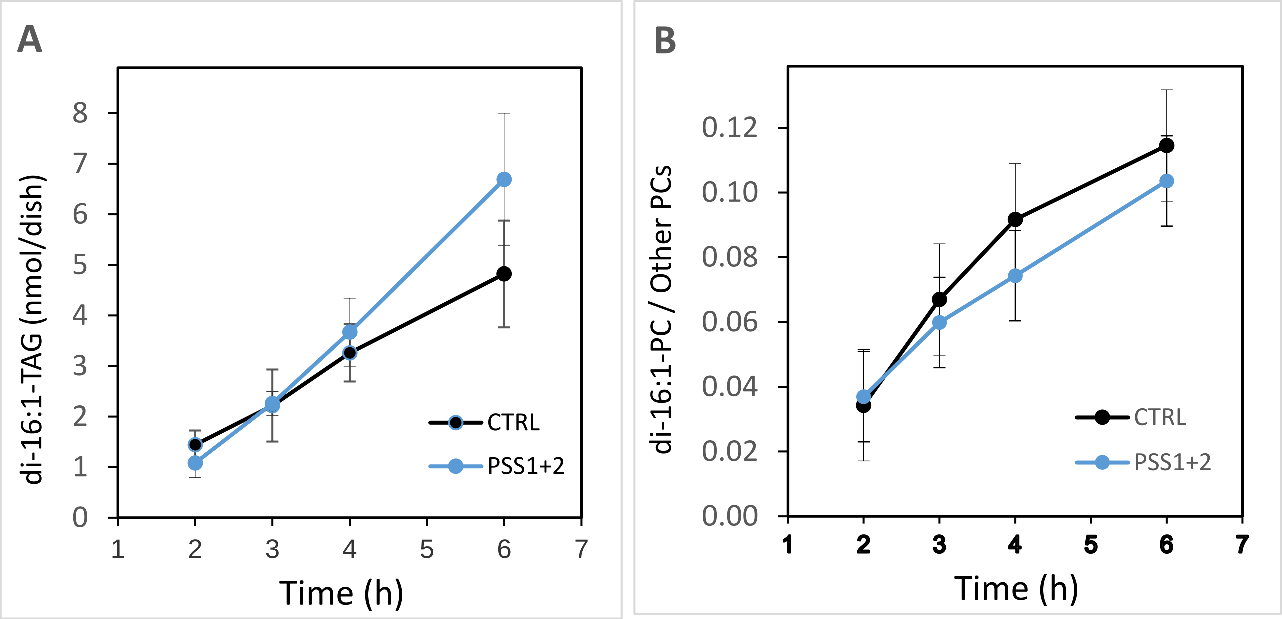


**Figure S5.** *Effect PSS1 + PSS2 knock down on the conversion of di-16:1-PS to di-16:1-TAG (A) and di-16:1-PC (B).* See Materials and Methods for details. The data are mean ± S.D. (*n* = 3). Black line = cells treated with control siRNA; Blue line = cells treated with PSS1 and PSS2 siRNA.

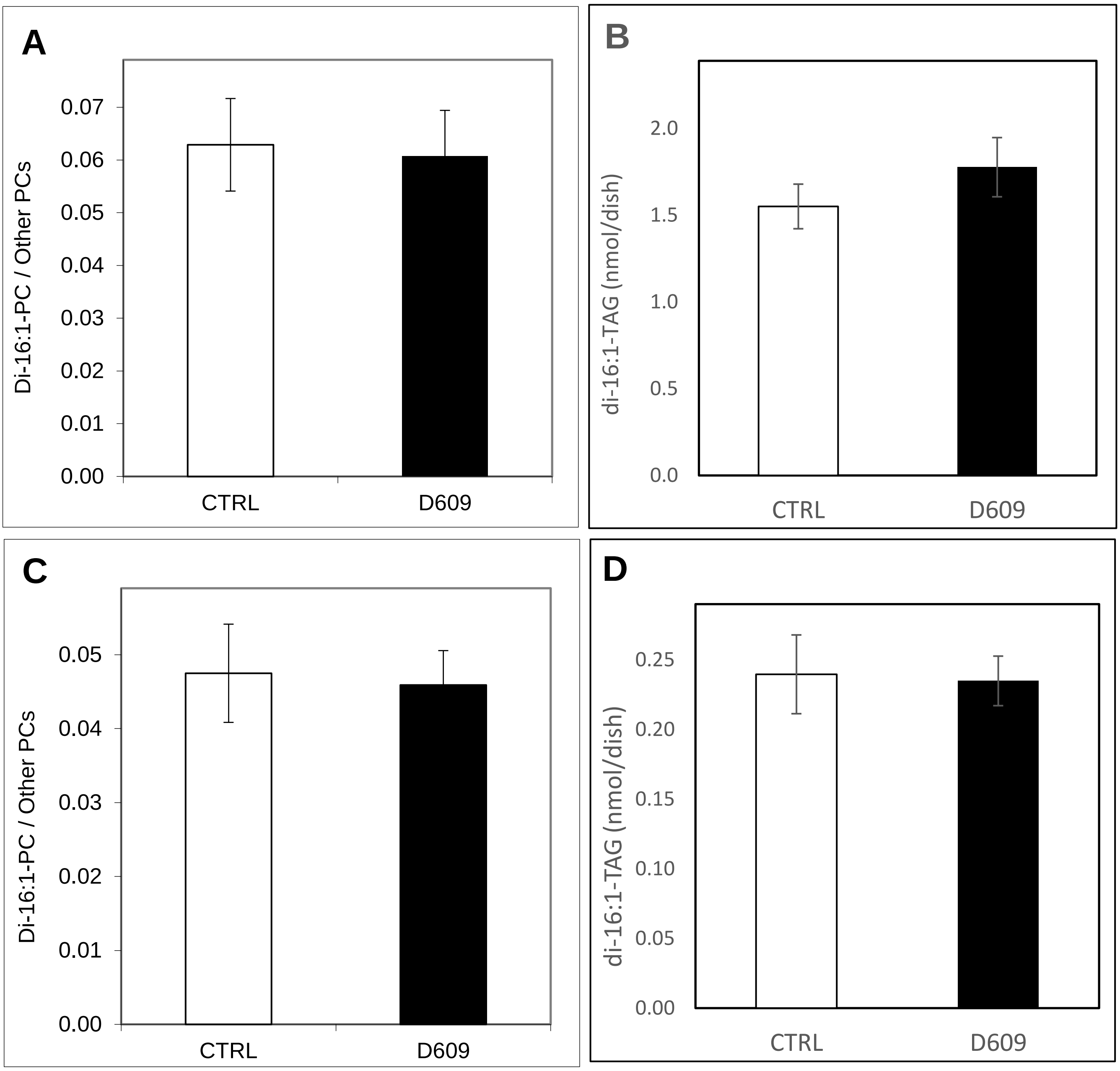


**Figure S6.** *Effect D609 on the conversion of di-16:1-PS (A,B) and di-16:1-PI (C,D) to di-16:1-PC (A, C) and di-16:1-TAG (B,D).* For details see legend of Fig. 2**.** Data are means ±S.D. (n = 3).

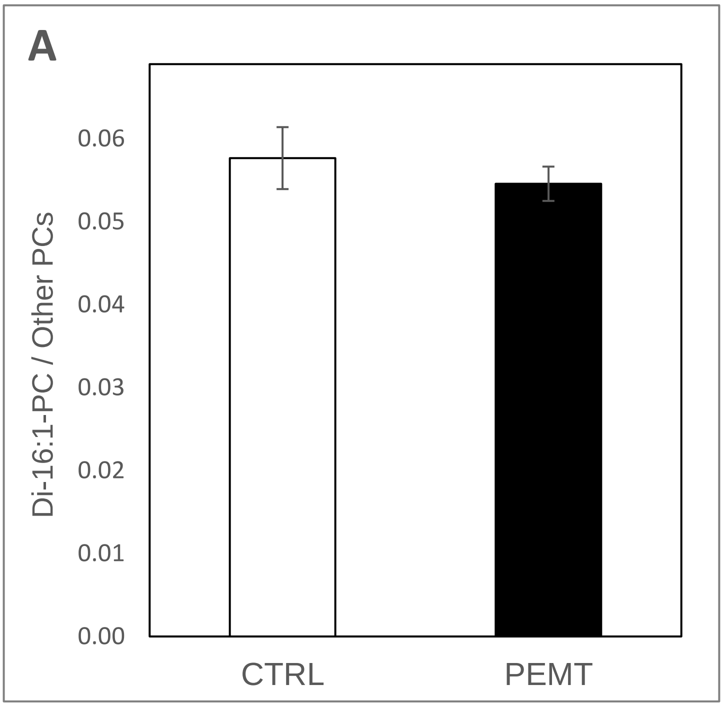


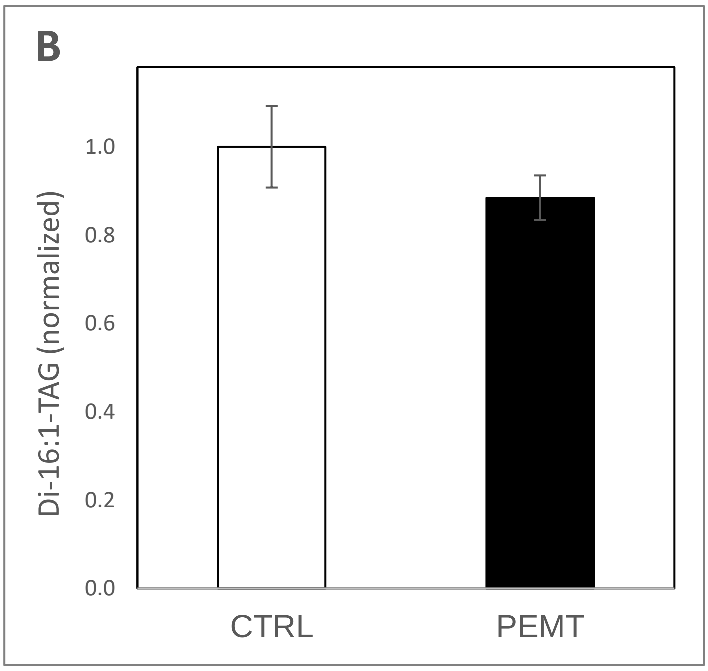


**Figure S7.** *Effect PEMT knock down on the conversion of di-16:1-PE to di-16:1-PC (A) or –TAG (B) in HeLa cells.* See Experimental Protocols for details. The data are mean ± S.D. (*n* = 3).

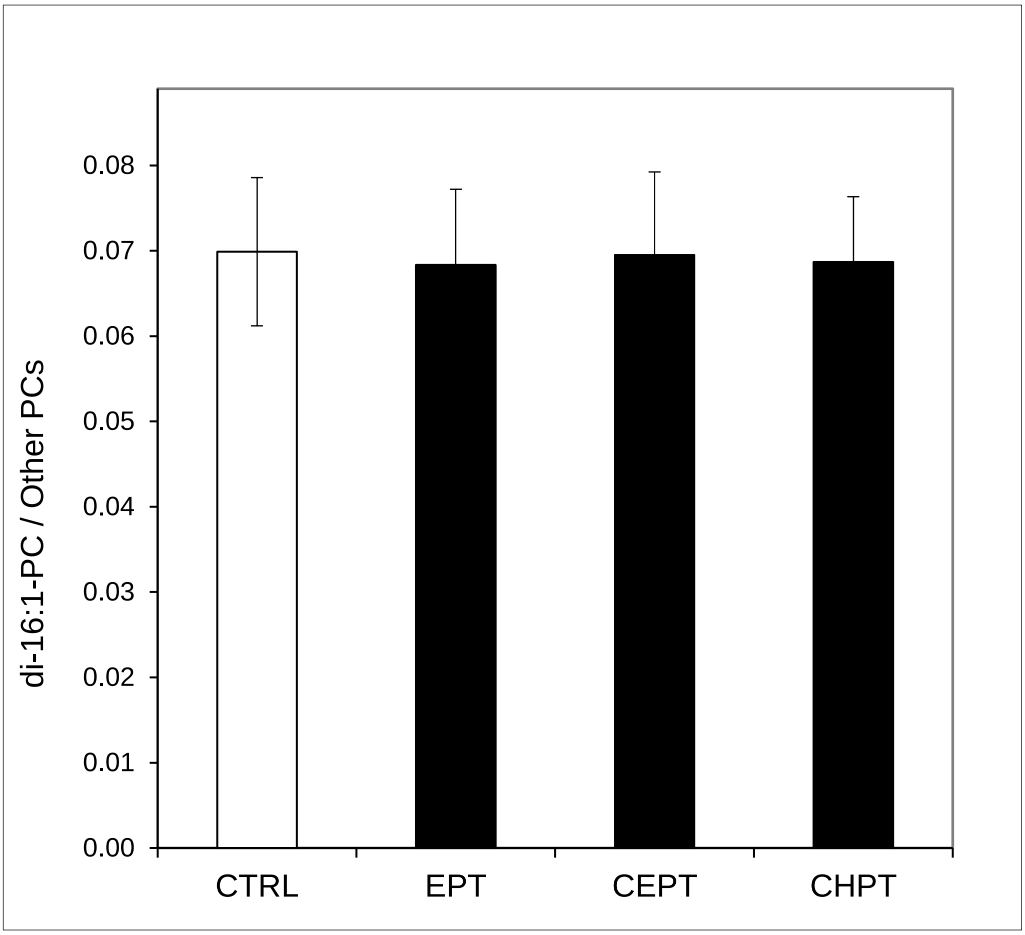


**Figure S8.** *Effect CEPT, EPT or CHPT knock down on the conversion of di-16:1-PE to di-16:1-PC (A) in HeLa cells.* See Materials and Methods for details. The data are mean ± S.D. (*n* = 3).

### Supplementary Tables

**Table S1.** ***siRNAs used in this study.***

| *Target gene* | *siRNA ID* | *Provider* |
|---|---|---|
| CEPT1 | SI00343770 | Qiagen |
| CHPT1 | SI04333728 | Qiagen |
| EPT1 | SI03131345 | Qiagen |
| PEMT | s195029 | Thermo Fisher (Ambion) |
| PLCB1 | SI02781184 | Qiagen |
| PLCB3 | SI00018781 | Qiagen |
| PLCD3 | s41419 | Thermo Fisher (Ambion) |
| PLCE1 | SI05056471 | Qiagen |
| PLCG1 | SI00041181 | Qiagen |
| PLCH2 | SI03066588 | Qiagen |
| PLCL2 | SI00686196 | Qiagen |
| PLD2 | SI00041237 | Qiagen |
| PLD3 | SI04238346 | Qiagen |
| PLD6 | SI00486024 | Qiagen |
| PTDSS1 | SI04169984 | Qiagen |
| PTDSS2 | SI104255965 | Qiagen |
| PLPPI | s16380 | Thermo Fisher (Ambion) |
| PLPP2 | s16383 | Thermo Fisher (Ambion) |
| PLPP3 | s16386 | Thermo Fisher (Ambion) |
| SGMS1 | SI03234917 | Qiagen |
| SGMS2 | SI02758182 | Qiagen |

***Table S2. Efficiency of siRNA mediated mRNA depletion.***

| siRNA | nM | mRNA expression[b] (% of CTRL ±SD) | Control genes[a] |
|---|---|---|---|
| CEPT1 | 2 | 11 ± 1.5 (n=3) | HMBS, SDHA |
| CHPT1 | 2 | 12 ± 3.1 (n=3) | SDHA, TBP |
| EPT1 | 2 | 9 ± 2.1 (n=3) | RPL13A, TBP |
| PEMT | 2 | 5 ± 1.4 (n=2) | HPRT1, SDHA |
| PLCB1 | 5 | 18 ± 0 (n=2) | HMBS, HPRT1 |
| PLCB3 | 2 | 16 ± 5.5 (n=3) | HPRT1, SDHA |
| PLCD3 | 5 | 28 ± 1.4 (n=2) | HPRT1, SDHA |
| PLCE1 | 2 | 27 ± 2.5 (n=3) | HMBS, TBP |
| PLCG1 | 5 | 9 ± 4.2 (n=2) | HPRT1, TBP |
| PLCH2 | 10 | 15 ± 4.0 (n=3) | HMBS, RPL13A |
| PLCL2 | 2 | 16 ± 4.9 (n=2) | HMBS, RPL13A |
| PLD2 | 5 | 25 ± 2.8 (n=2) | HMBS, SDHA |
| PLD3 | 5 | 6 ± 3.5 (n=2) | HMBS, TBP |
| PLD6 | 2 | 23 ± 0 (n=2) | HMBS, TBP |
| PTDSS1+PTDSS2 | 2+2 | 6 ± 1.5, 15 ± 2.0 (n=3) | HMBS, HPRT1, RPL13A |
| SGMS1 | 5 | 16 ± 1.4 (n=2) | HPRT1, TBP |
| SGMS2 | 5 | 21 ± 2.8 (n=2) | HPRT1, TBP |

[a] Control genes are selected based on target stability value assessed by CFX Manager Software.
[b]For PLPPI, PLPP2 and PLPP3 siRNAs confirmed by the manufacturer were used.

***Table S3. qPCR primers used in this study***. The primers were either designed using the NCBI Primer-BLAST software or the sequences was obtained from publications.

| Gene | Forward and reverse sequences 5' – 3' | |
|---|---|---|
| CEPT1 | TTGTGCGCACTGGCAAACGTA<br>GGTGGTCCTCCAATCACTGCCA | |
| CHPT1 | AGGGGAGCTCTTTGACCATGGCT<br>GGATAAGTTCCTAAGCGAGCGGCA | |
| EPT1 | GCCTTGTAGCCGGGAGTCGC<br>AAGTGGATTGGTATCCACAGCACTG | |
| PEMT | CCTGGATCCCAGCTTTGTGG<br>GGGTCTTGTGTTCCCATCGT | |
| PLCB1 | AGCCGCTTTGGAAAAGTCCGC<br>TCCAGAGCAGCGAGGTCTTGCT | |
| PLCB3 | ATGGGACGAGGAGAAGCTGATGAC<br>GGGAGGCGTTCTGAGCCAGGAT | |
| PLCD3 | CGCCATCGCAGCACATCTTCTTCG<br>CAGGTTCTTGCGGCGGCCCTTG | |
| PLCE1 | CCCAACTTGGCTGCCGGGAC<br>TGCCGCCTTTGATCCGCCC | |
| PLCG1 | GCTCAGGCTACTTTCCCAGTA<br>AGGGACTGAGGTCACCATTCT | |
| PLCH2 | AAGCTCCCAGCCAACATCAG<br>ACACGCTTTCGATTGGTGGA | |
| PLCL2 | AAGAATGCCCAGCCCCCTATACGG<br>AAAGCGGGGAGACACCGCCAA | |
| PLD2 | GTCTGCCAGGGATGACGGCG<br>AAACGGGTGCATCCGGTCGG | |
| PLD3 | GCTGACCCATGGCGTCCTGC<br>CGAGCCAGGCAGCTGCAGTT | |
| PLD6 | GGCCCTCAACGGCTCGCAAA<br>TGGATGGCTTGCGTGGTCCAG | |
| PTDSS1 | GGTGGCATCACAGCTCCCACA<br>ACCCAGCATTGTGTTCCTACGCG | |
| PTDSS2 | TGGCTTTGTTCCCGCGCACT<br>GTTGGGCAGCTGGTGCTCCA | |
| SGMS1 | TAACCGGGTGCAGTGGGCCT<br>ACCGATACAGGTACAGCGTGCCA | |
| GMS2 | CGGTCACACGGTTACGCTGACAC<br>GCAGATGATCCCGGCAGCACTC | |
| HMBS | TGCAACGGCGGAAGAAAA<br>ACGAGGCTTTCAATGTTGCC | Ref. 1 |
| HPRT1 | TGACACTGGCAAAACAATG<br>GGTCCTTTTCACCAGCAAGCT | Ref. 1 |
| RPL13A | AAAAAGCGGATGGTGGTTC<br>CTTCCGGTAGTGGATCTTGG | Ref. 2 |
| SDHA | TGGACCTGGTTGTCTTTGGT<br>AGTCGCAGTTCCGATGTTCT | Ref. 2 |
| TBP | TGCACAGGAGCCAAGAGTGAA<br>CACATCACAGCTCCCCACCA | Ref. 2 |

*Reference 1:* Cicinnati, V. R., Shen, Q., Sotiropoulos, G. C., Radtke, A., Gerken, G., and Beckebaum, S. (2008) Validation of putative reference genes for gene expression studies in human hepatocellular carcinoma using real-time quantitative RT-PCR. BMC Cancer 8, 350
*Reference 2:* Pombo-Suarez M, Calaza M, Gomez-Reino JJ, Gonzalez A. (2008) Reference genes for normalization of gene expression studies in human osteoarthritic articular cartilage. BMC Mol Biol. 2008 29, 9:17.
